# Comparative survey of the relative impact of mRNA features on local ribosome profiling read density

**DOI:** 10.1101/018762

**Authors:** Patrick BF O’Connor, Dmitry E. Andreev, Pavel V. Baranov

## Abstract

Ribosome profiling (Ribo-seq), a promising technology for exploring ribosome decoding rates, is characterized by the presence of infrequent high peaks in ribosome footprint density and by long alignment gaps. Here, to reduce the impact of the data heterogeneity we introduce a simple normalization method, Ribo-seq Unit Step Transformation (RUST). RUST is robust and outperforms other normalization techniques in the presence of heterogeneous noise. We illustrate how RUST can be used for identifying mRNA sequence features that affect ribosome footprint densities globally. We show that a few parameters extracted with RUST are sufficient for predicting experimental densities with high accuracy. Importantly the application of RUST to 30 publicly available Ribo-seq datasets revealed a substantial variation in sequence determinants of ribosome footprint frequencies, questioning the reliability of Ribo-seq as an accurate representation of local ribosome densities without prior quality control. This emphasizes our incomplete understanding of how protocol parameters affect ribosome footprint densities.

## Introduction

The advent of ribosomal profiling (ribo-seq) has provided the research community with a technique that enables the characterization of the cellular translatome (the translated fraction of the transcriptome). It is based on arresting translating ribosomes and capturing the short mRNA fragments within the ribosome that are protected from nuclease cleavage. The high throughput sequencing of these fragments provides information on the mRNA locations of elongating ribosomes and thereby generates a quantitative measure of ribosome density across each transcript. Accordingly, ribosome profiling data contain information that could be used to infer the properties that affect ribosome decoding (or elongation) rates. Unsurprisingly, a large number of studies analyzing ribosome profiling data for this purpose have been published recently^1-21^.

There is a considerable discordance among some of the findings in these works that is unlikely to be wholly caused by differences in the biological systems used. It may also be attributed to the computational methods used for estimating local decoding rates which are often based on elaborate models of translation that use certain assumptions regarding the process. The abstraction required for modelling necessitates the generalization of the process across all mRNAs, although we are aware of numerous special cases^22^. Even if the generalized models provide an accurate representation of the physical process of translation in the cell, they do not model the ribosome profiling technique itself, which may introduce various technical artefacts. Oft-cited potential artefacts include the methods used to arrest ribosomes (the result is affected by the choice^8,23^ and the timing^7,21,24^ of antibiotic treatment), the sequence preferences of enzymes involved in the library generation^1,25^ and the quality of alignment. These artefacts may distort the output and it may not be easy to disentangle their effects in the presence of biologically functional and sporadic alterations in translation.

Ribosome profiling data are characterized by high heterogeneity caused by alignment gaps and sporadic high density peaks due to technical artefacts and ribosome pauses^4,26^. These fluctuations, even if caused by genuine ribosome pauses, are thought to negatively impact the ability of some methods to accurately characterize factors that influence ribosome read density globally. With this rationale we developed a data smoothing method, that we term RUST (Ribo-seq Unit Step Transformation). We first demonstrate that RUST is resistant to the presence of heterogeneous noise using simulated data and outperforms other normalization techniques in reducing data variance. Then we analyze real data from 30 publicly available ribosome profiling datasets obtained using samples (cells or tissues) from human^14,27-39^, mice^7,37,40-42^ and yeast^1,6,8,12,43-45^.

We show that a few parameters extracted with RUST are sufficient to predict experimental footprint densities with high accuracy. This suggests that RUST noise resistance allows accurate quantitative assessments of the global impact of mRNA sequence characteristics on the composition of footprint libraries.

The comparison of RUST parameters among different datasets revealed a considerable discordance in the relative impact of the sequence factors determining frequencies of ribosome footprints in the libraries. This most likely can be attributed to the differences in experimental protocols, suggesting that the variance in the data, rather than in the analytical approaches used is responsible for the current contradictions regarding the sequence determinants of the decoding rates.

## Results

### Ribo-seq Unit Step Transformation (RUST)

The probability of finding a ribosome decoding a particular codon of an mRNA (and by extension the expected number of corresponding ribo-seq reads in a library) depends on three variables: the mRNA expression level, the translation initiation rate for the corresponding open reading frame (ORF) and the time that the ribosome spends at that codon (dwell time). The latter (as an invert) is usually described as a codon elongation rate or a codon decoding rate. Estimating the true decoding rates with ribo-seq is made difficult by the absence of precise measurements of initiation rates. Therefore, studies (including this one) using ribo-seq for this type of analysis typically attempt to measure the relative dwell time of codons instead of the actual dwell time. A frequent and intuitive approach is the normalization of the local ribo-seq signal by the average signal across the coding region^4,9^. This approach has been described as conventional^4^ and we will refer to it as CN for conventional normalization. It is based on the reasonable assumption that the transcript expression levels and ORF initiation rates are the same for all codons from that ORF. CN is perceived to have two major shortcomings: it is expected to be very sensitive to the high density peaks which frequently occur due to functional ribosome pauses^4^ (Fig. 1a) and it is typically applied only to the transcripts with high ribosome coverage, as the relative impact of a single read alignment on CN is excessive with sparse profile data (Fig. 1a).

**Figure 1.**
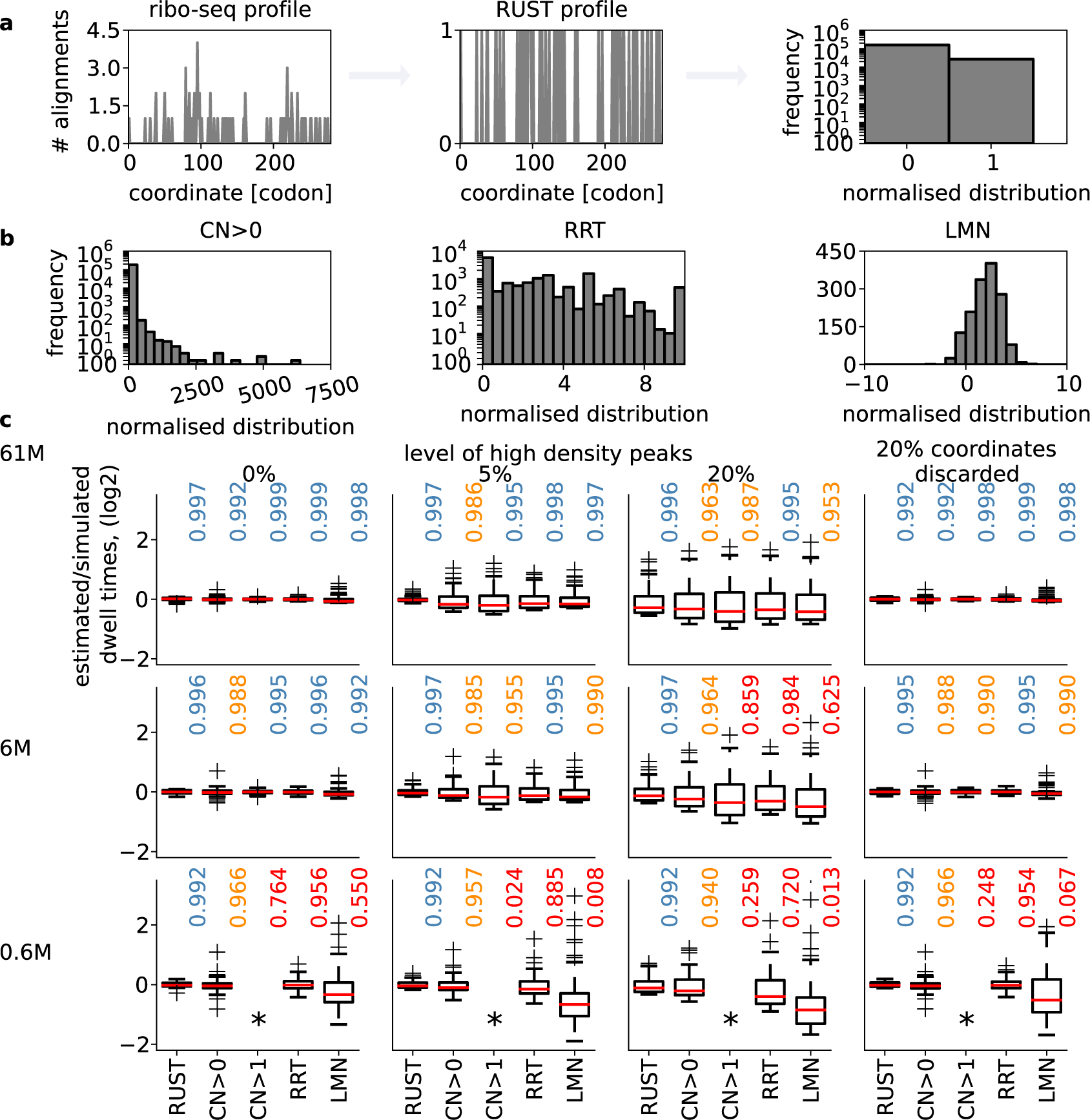
Comparison of ribosome profiling normalization approaches. **(a)** A stylized footprint density profile for *MTIF3* gene transcript from ‘Andreev’ dataset (left) is transformed into a binary function with RUST (center). Each sequence feature, such as AAA codon in the case shown, could be characterized by its frequency as 1 or 0 (right). **(b)** The distributions of normalized codon densities for all AAA codons in ‘Andreev’ dataset using different approaches, conventional normalization CN (left), ribosome residence time, RRT (top right) and logarithmic mean normalization LMN (right). Note that due to intrinsic differences the scale of possible normalized densities (axes X) varies among the methods and that due to the selection criteria of each approach the number of datapoints used (axes Y) is also variable. **(c)** Performance of five normalization approaches (RUST, CN of transcripts with average gene density >1/nucleotide (CN>1) and CN of all expressed transcripts (CN>0), LMN and RRT) at estimating codon dwell times The box plots show the distribution of log values of the estimated/simulated dwell times for all 61 codons. The deviations of these values from 0 occur due to under or overestimation of simulated dwell times. The better methods are those that have distributions with a smaller variance. Each subpanel represents a specific scenario. The simulation scenarios differ by coverage that reduces from top to the bottom and the level of noise modelled as high peaks of density that increases from left to right, except for the right-most column where noise is modelled as missing data at 20% of the coordinates. Asterisks used to indicate insufficient data for CN>1.

Various approaches have been tried to reduce the impact of density outliers (Fig. 1b). Dana and Tuller removed atypical densities based on expected distribution of densities^4^. Artieri and Fraser used logarithmic mean instead of the arithmetic mean to produce a “corrected ribo coverage”^1^ (Fig. 1b). Gardin et al.^6^ developed an intricate approach for calculating a statistics that they called “ribosome residence time (RRT)”. The approach involves CN like sampling, but only from specific segments of RNA that satisfy certain sequence and coverage requirements^6^ (Fig. 1b). Pop et al. introduced a sophisticated model that is based on the assumption that the ribosome footprint density profile must satisfy flow conservation constraints, i.e. the translation is at steady state and that all ribosomes translated the entire coding region^12^. While flow conservation constraints may be true for the ribosome densities, they may not hold for footprint densities because of technical artefacts such as sequencing biases and misalignments.

We reasoned that a practical approach for the analysis of ribosome profiling data should be (i) simple; (ii) robust to the presence of heterogeneous noise; (iii) able to use all available data (i.e. no restriction to genes with high read coverage) and (iv) be able to produce statistics that would allow accurate prediction of experimental densities. With this in mind we developed a procedure that we term Ribo-seq Unit Step Transformation (RUST) where the ribosome footprint densities (the number of reads corresponding to the position of the A-site codon) are converted into a binary step unit function (also known as Heaviside step function). Individual codons are given a score of 1 or 0 depending on whether the footprint density at these codons exceeds the average for the corresponding ORF (Fig. 1a and Supplementary Fig. 1). In addition to codons, the procedure could be applied to any other potential determinant of read density such as individual nucleotides, encoded amino acids, their combinations as well as their properties, such as a charge or hydrophobicity of encoded peptides or free energy of RNA secondary structures. The average RUST value for each putative determinant of decoding rates may be compared to the expected RUST value to measure its effect, see Methods. As a result of the transformation the impact of every site has a small influence on the final RUST value. The value is influenced primarily by the consistent presence of reads at numerous sites. For example, no differentiation is made between a stall site where the ribosome density just exceeds the average to one where the average is grossly exceeded. For the details of transformation, see Methods and the RUST pipeline in Supplementary Figure 1.

### Evaluation of normalization methods with simulated data

In order to evaluate RUST performance, we tested its ability to estimate decoding rates from simulated data. We simulated the data under a simplifying assumption that the local decoding rates depend only on the identity of a codon in the A-site. To simulate the data we used real transcript sequences and experimental distribution of footprints per transcript, but modeled the distribution of footprints within a transcript by specifying the dwell time of each of 61 codons and introducing different levels of heterogeneous noise (see Methods and below for how the noise was simulated). We compared its performance to the RRT approach, the CN method and to a logarithmic mean normalization (LMN) similar to that obtained with the “corrected ribo coverage” (see Methods). Unlike in the original approach in LMN ribo-seq density is not normalized by the mRNA-seq density. The CN method was used in two modes with filtering requiring a minimal coverage threshold (average transcript footprint density of >1 read/nucleotide) CN>1, and without any threshold, CN>0. The parameters of the simulation were selected either to produce data similar to the experimental data or the data with reduced quality (see Methods). For example, the sequencing depth was either equal or lower than what has been obtained with actual data.

Figure 1c compares the performance of the five methods for three different simulated sets of data with different sequencing depth and levels of noise simulated as sporadic high density peaks (3× the value of the highest footprint density for the original simulated profile) or as a loss of density that could arise, for example due to removal of ambiguous mappings. For these simulations the relative time that ribosome dwell at each of 61 codons *t_c_* was pre-set (see Methods) and the normalization approaches were compared in their ability to accurately detect codon dwell times (*t_c_*) from the simulated data. The estimated-to-simulated dwell time log ratios were obtained for 61 codons. We assessed the performance of each method by showing the distribution obtained using box plots. For accurate methods the values for each codon should be zero, i.e. the observed and simulated values should be the same. We also provide the coefficient of determination, R^2^, between the estimated and simulated dwell times as a measure of the normalization approaches accuracy, with values closer to one indicating better accuracy. We find that all approaches estimate relative *t_c_* values very accurately in the absence of noise provided that coverage is high. However, in the presence of noise or under reduced coverage the performance worsens. In this regard RUST appears to be the most resilient to the reduced coverage and both types of noise. While its ability to accurately predict simulated relative dwell times drops under high levels of noise, the combined inferred values still correlate remarkably well with the simulated values (Fig. 1c).

We conjectured that the accuracy of the normalization approaches may depend on codon specific properties, such as a relationship between codon usage and dwell times. Therefore, we simulated the data under three different sets of *t_c_* parameters. In the first two simulations the range of *t_c_* values were set to rank-correlate with the codon usage (see Methods and Supplementary Fig. 2), i.e. the lowest *t_c_* was set for the rarest codon and the highest *t_c_* for the most abundant codon. In one set the *t_c_* range spans one order of magnitude and in the other, two orders of magnitude. In the third set, the *t_c_* parameters were set to negatively correlate with the codon usage. For the scenario where the range of decoding rates is increased to span two orders of magnitude (Supplementary Fig. 2, middle and bottom plots) the effect of noise on the accuracy of *t_c_* inference is similar.

**Figure 2.**
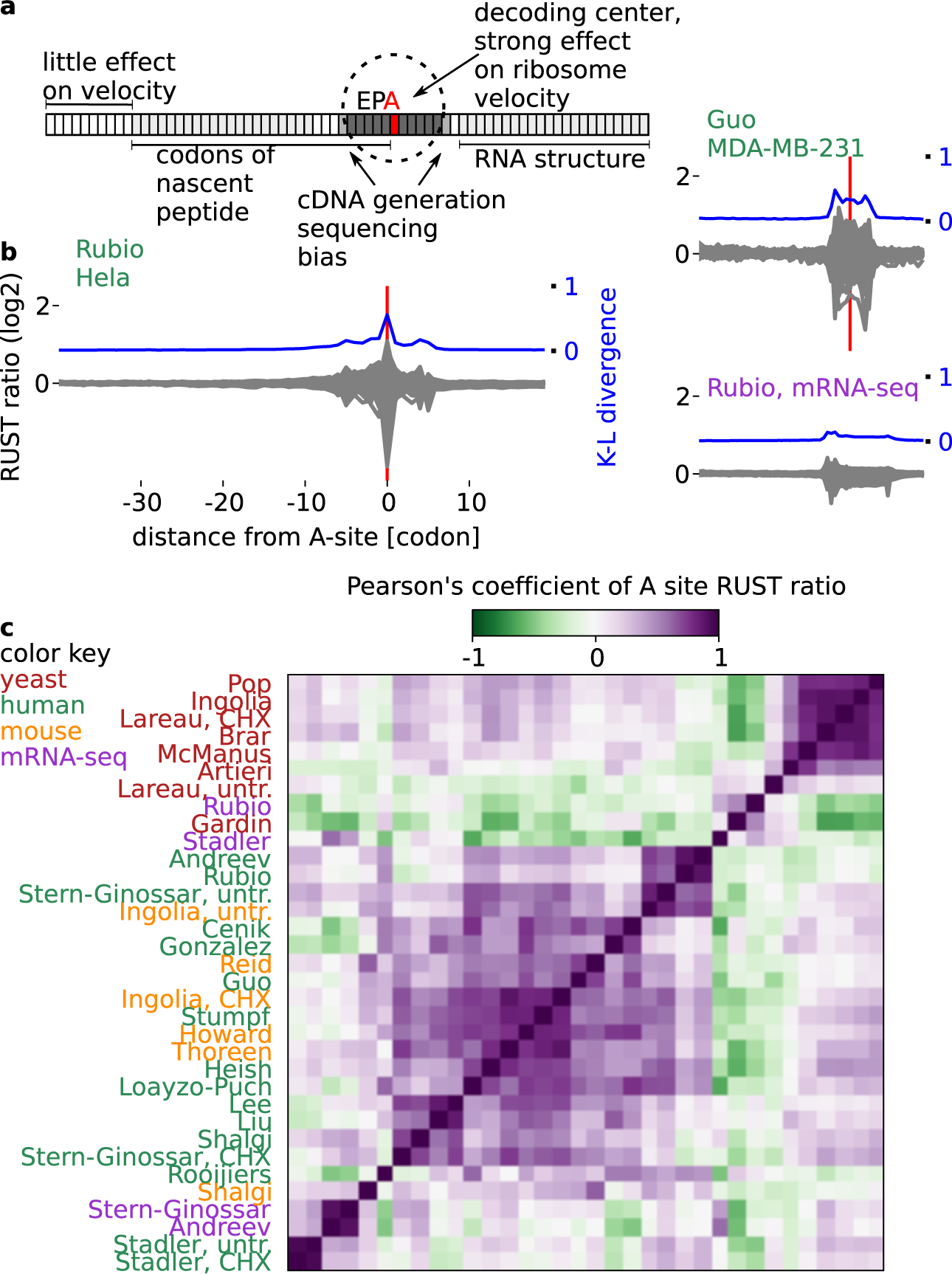
Evaluation of ribo-seq datasets with RUST. (a) Anatomy of the ribosome footprint displaying position-specific mRNA sequence influence on ribo-seq read density. (b) RUST codon metafootprint profiles of selected ribo-seq and mRNA-seq datasets used in this study. The individual RUST ratio values of 61 sense codons across the mRNA are displayed. The resulting grey area is a superposition of each 61 curves. The corresponding Kullback-Leibler divergence (K-L) is shown in blue. The protocol details for each dataset are summarized in Table 1.(c) Heatmap displaying the pairwise similarity of codon RUST ratios at the A-site, as measured by the Pearson’s correlation, for ribo-seq datasets of human (green), yeast (red) and mouse (orange). Also included are human mRNA-seq data (violet). The datasets are indexed by the name of the first author. The clustering was done with Scipy using the “Euclidean” distance metric with “single” linkage.

Interestingly, in all simulations (Fig. 1c and Supplementary Fig. 2) the logarithm ratios between estimated and simulated values are not uniform among 61 codons, i.e. the estimations are not equally accurate for each codon. The estimated relative dwell times of quickly decoded codons were found to be consistently overestimated by all methods tested, i.e. inferred as slower. This is more acute when the decoding rates span 2 orders of magnitude but is also observed even when the decoding rates span 1 order of magnitude (Supplementary Fig. 2, top plots). We also found that the R^2^ values were consistently lower when the codon usage negatively correlated with the simulated dwell time than when they were positively correlated (Supplementary Fig. 2, middle and bottom plots). However, the difference is small suggesting that relationship between the codon usage and decoding rate appears to have a relatively minor influence on the correct estimation of the relative dwell time.

Counterintuitively in most simulations CN>0 performs similar or even better than CN>1 and the LMN was found to be inferior to both CN normalizations. Under almost all scenarios tested RUST was found to outperform other normalization techniques in the presence of noise.

### The impact of technical biases varies among datasets

The velocity of a ribosome could be influenced by the sequence of mRNA in several ways (outlined in the scheme in Fig. 2a). Codons in the E-, P- and A-sites of the ribosome determine the identity of corresponding tRNAs (and amino acids) inside the ribosome. The mRNA sequence in the cavity between subunits could affect ribosome movement by directly interacting with its components. In addition, the sequence upstream of the A-site codon (up to 90 nucleotides) could influence the progressive movement of the ribosome through the interactions between the peptide it encodes and ribosome peptide tunnel. Lastly, the sequence downstream of the ribosome could alter its velocity through the formation of stable RNA secondary structures^46,47^ or the presence of RNA-protein complexes.

In addition to these intrinsic factors affecting ribosome velocities, there are technical factors that influence the distribution of sequencing reads in ribo-seq datasets. First, the drugs used to block elongating ribosomes could act on ribosomes only at a specific conformation^8^ or could also alter their distribution along mRNAs^23,24^. Second, various enzymes used for cleaving mRNA, for generating cDNA libraries and for their sequencing exhibit sequence specificity especially at the boundaries of ribosome footprints^1^. Third, the accuracy of alignment step depends on the existence of paralogs and transcript sequence complexity and the way how ambiguous alignments are treated. Fourth, the occurrence of alternative splicing, ribosome drop-off, ribosome stacking and alternative translation initiation may all affect the distribution of reads across individual transcripts.

To analyze how sequence of mRNA effect density of footprints in different locations relative to the A-site in experimental data we used an approach similar to the one used by Artieri and Fraser^1^. We calculated observed-to-expected RUST ratios for each codon position within a window of 60 codons (see Methods and Supplementary Fig. 3). This window encompasses the ribosome protected fragment (codons -5 to +5), the region encoding the nascent peptide (codons -30 to 0) and the region downstream of the ribosome (+5 to +20), where zero coordinate corresponds to the A-site codon. To measure the contribution of local mRNA positions to the density of footprints correspondingly derived from a ribosome decoding a particular codon, we measured the relative entropy at each position using the Kulback-Leibler (K-L) divergence.

**Figure 3.**
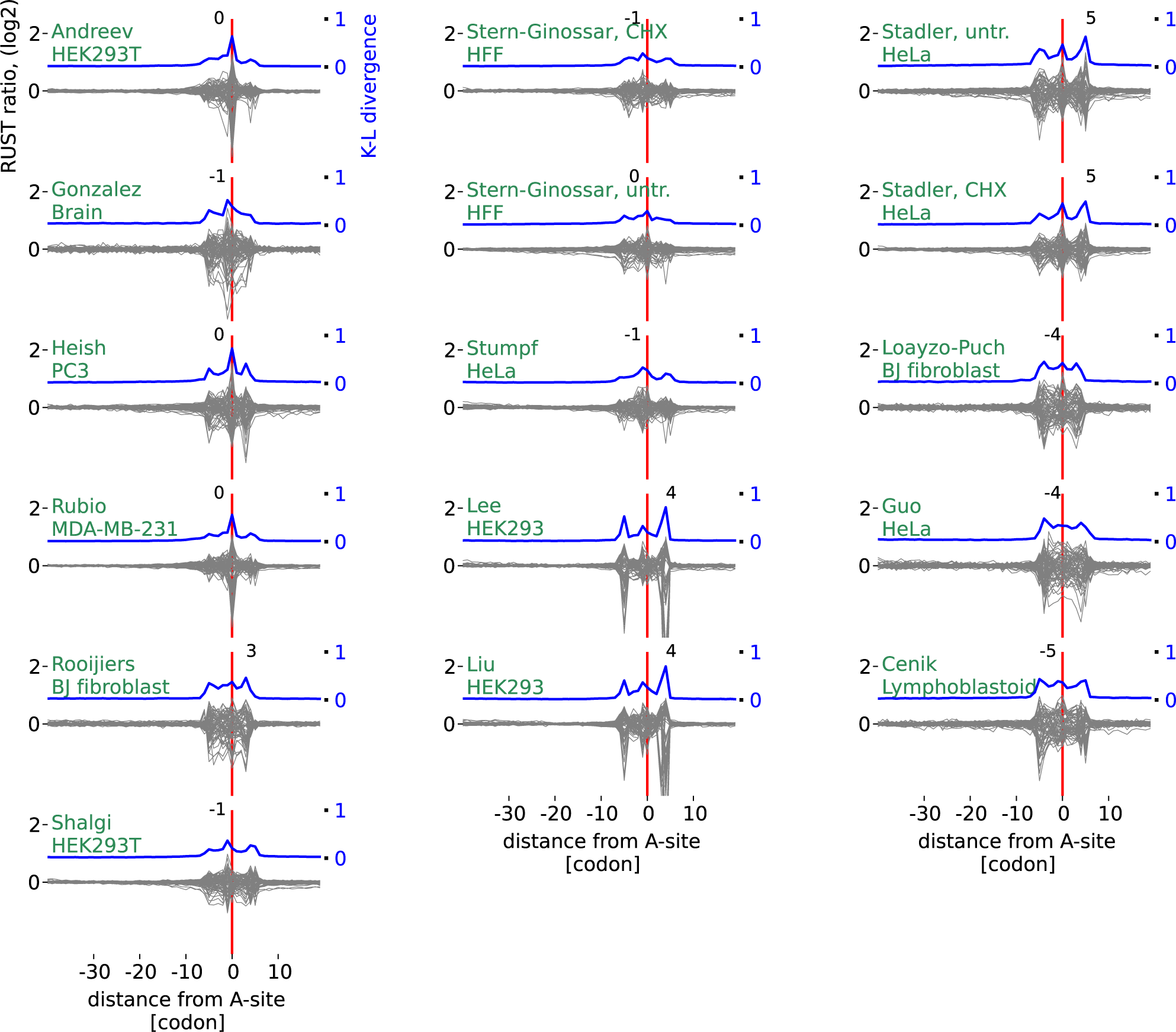
RUST metafootprint profiles of the 16 human ribo-seq datasets. Datasets are indexed by the name of the first author followed by drug treatment and source, see Table 1 for more details. The Kulback-Leibler (K-L) divergence is shown in blue, the coordinates of K-L maximum are indicated above the peak in each plot. Zero coordinate corresponds to the inferred position of the A-site and is marked with a red line, coordinates are in codons. See Supplementary Fig. 4 for non-human studies.

Figure 2b shows the relative entropy and normalized observed-to-expected RUST ratios *ro/re* (see Methods) for each individual codon for two of the ribosomal profiling datasets explored in this work. By analogy with metagene profiles we refer to the plots of *ro/re* RUST ratios as metafootprint profiles. The areas of reduced entropy (increased K-L divergence) are mostly contained within a window of 10 codons upstream and downstream of the A-site, approximately matching to the position of the actual ribosome footprint. In almost all cases three local K-L maxima are observed, one corresponds to the decoding center (Fig. 2b), the other two maxima roughly correspond to the 5’ and 3’ ends of ribosome footprints. The same procedure carried out on mRNA-seq libraries reveals decreased entropy in the same area with two maxima corresponding to the mRNA fragment ends (Fig. 2b). This suggests that the main contributing factors to footprint frequency at the corresponding location are the identity of the codons in the A- and/or P-sites and the sequence-specificity of the enzymes used during library construction. The metafootprint analysis for all human studies explored in this study are available in Figure 3, see Supplementary Figure 3 for non-human studies and mRNA-seq controls. The degree of variation in the relative impact of these factors among different datasets is surprising. In some of the ribo-seq datasets, the density of footprints depends on the identity of the codon at the ends of footprint more than on the identity of the codon in the A- or P-sites. This is suggestive of a high level of sequencing biases introduced during the cDNA library generation in some of the tested datasets.

Figure 2c shows a heatmap produced as a result of pairwise comparison of observed-to-expected RUST ratios for the 61 codons when they are located in the A-site. Most apparent is the high reproducibility for most ribosomal profiling datasets produced in yeast under cycloheximide pretreatment (Fig 2c, Supplementary Fig. 5). The comparison of the protocol conditions (Table 1) points to the consistency in the protocols used in these studies. The variance across the datasets obtained from mammalian sources is more substantial as are the differences in the protocols (Table 1). We found that variance in RUST ratios of nonsynonymous codons is greater than that of synonymous codons. In other words, the identity of decoded amino acid has a greater influence on read density than the identity of the specific codon. ANOVA revealed that this was statistically significant in 28 of the 30 ribo-seq samples. We carried out similar analysis for mRNA-seq controls for codons located at the same distance from the 5’ end as the A-site codons in ribosome footprints. As expected, the degree of variation among all 61 codons was much smaller. However synonymous codons also exhibited statistically significant higher variation (Supplementary Fig. 6). This casts some doubts on biological relevance of this observation.

**Table 1.**
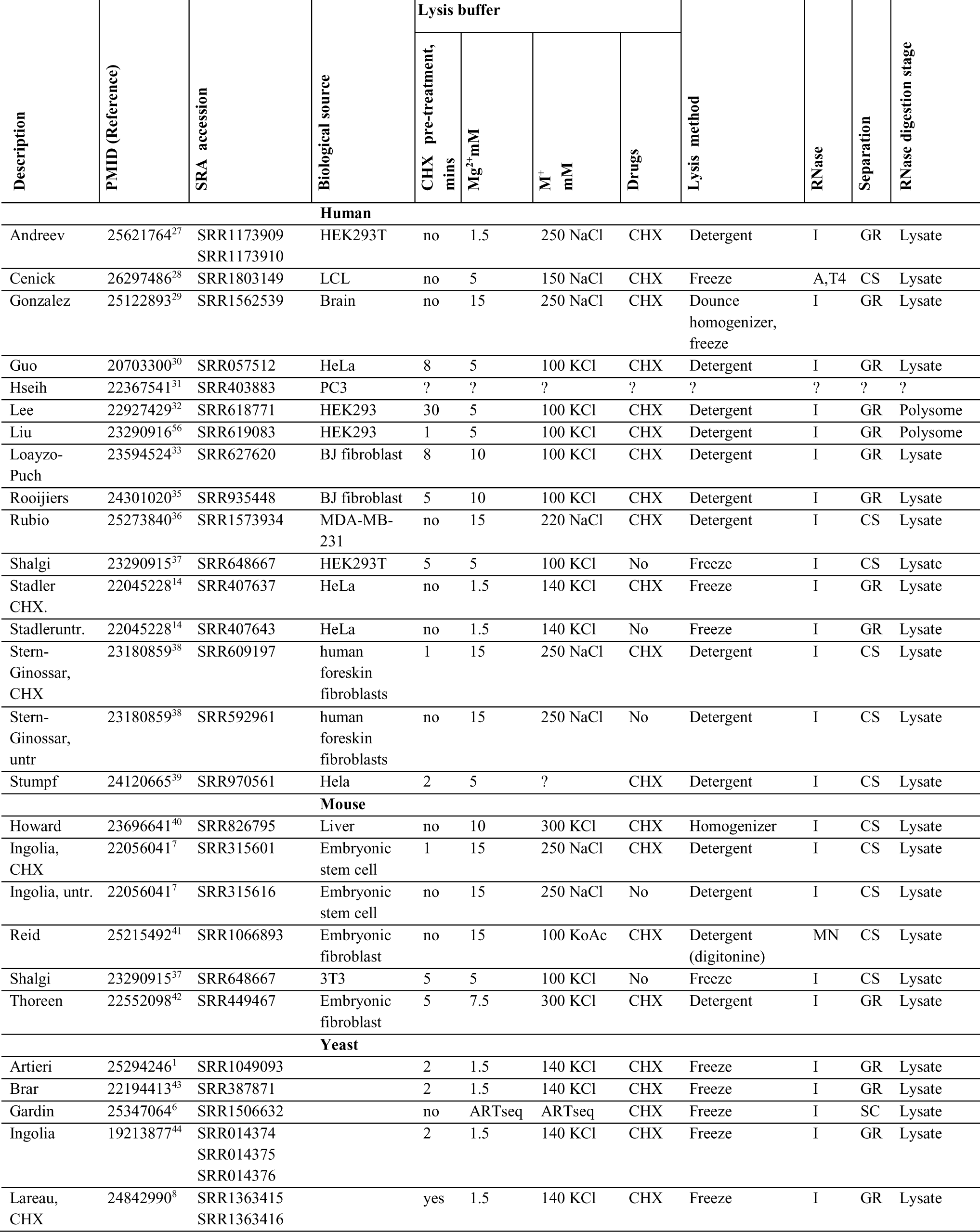
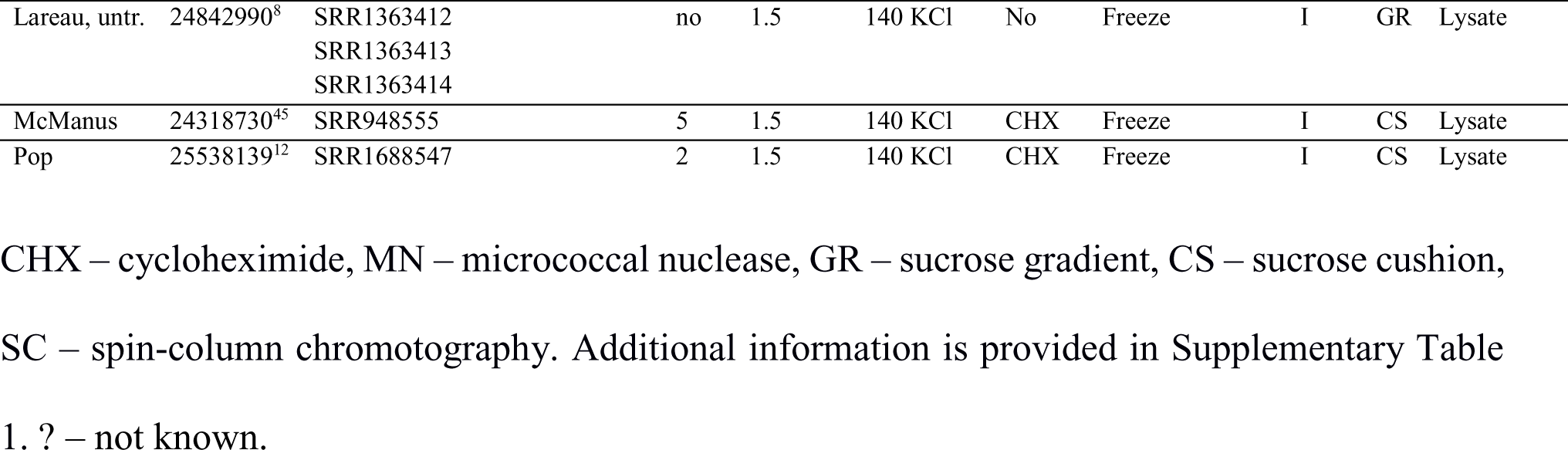
Ribosome profiling protocol conditions for the studies described in this work

Some of the studies produced the data with a change to a single parameter: the samples were either pretreated or not with cycloheximide before lysis ^7,8,14,38^. We found that ‘Stadler’^14^ datasets are similar for both types of treatments, while ‘Lareau’^8^, ‘Ingolia’^7^ and ‘Stern-Ginnoassar’^38^ are different (Fig. 2c). Supplementary Figure 7 provides the analysis of RUST ratios for ‘Lareau’ and ‘Ingolia’ datasets under both conditions, clearly indicating that cycloheximide substantially alters the distribution of footprints on mRNA. This is consistent with the observation that cycloheximide blocks ribosomes in a specific conformation and this ribosome arrest has certain codon preferences^16^. A more focused and detailed analysis of this phenomenon^23^ was published while this manuscript was in preparation.

Prior studies explored the effects of different antibiotic treatments in mammalian cells^7^ and in yeast^8,23,24^. The effect of buffer conditions on triplet periodicity was also explored to some extent^38,43^ as well as conditions of nuclease treatments^48^. We agree with a plea for standardization of ribosome protocols^25^, however, as recently argued^21^ it is clear that a more systematic study of protocol dependency of ribosome profiling data is needed for this.

### Influence of RNA secondary structure and nascent peptide

To illustrate RUST capacity at analyzing mRNA features that may affect ribosome velocities we chose three, ‘Andreev’^27^, ‘Rubio’^36^, ‘Pop’^12^. These datasets exhibit a low level of K-L divergence at the ends of the footprints and a high K-L divergence at the decoding center, suggesting low sequencing bias at the end of footprints. However, while these datasets are relatively free of sequencing artefacts, the distribution of footprints could still be skewed for other reasons discussed in the previous section and caution needs to be applied in the interpretation of the results described below.

To estimate the effect of RNA secondary structure we calculated the RUST ratios for RNA sequences that can form secondary structures at a particular free energy threshold as calculated with RNAfold^49^, see Methods. Supplementary Figure 8a shows the distribution of RUST ratios for RNA secondary structures predicted within 80 nucleotides window with different free energies. It can be seen that sequences predicted to contain stable structures are underrepresented (low RUST ratios) in windows that overlap with sequencing reads. This is observed for both ribo-seq and mRNA-seq reads and therefore is likely to be an artefact related to cDNA library generation and sequencing. This is not explained by a putative nucleotide bias. The distribution of individual nucleotides at the footprint location does not deviate significantly with the exception for the location of the decoding center (Supplementary Fig. 8b).

The RUST ratios for individual amino acids and dipeptides (Supplementary Fig. 9) do not reveal evidence of universal nascent peptide effect on ribosome velocity from the positions distant from the peptidyl transferase center. Although, such effects can be seen in individual datasets, e.g. strong influence of two Prolines in close proximity to the peptidyl transferase center in ‘Andreev’ dataset (Supplementary Fig. 9). Such nascent peptide interactions may also be facilitated by specific physicochemical properties of the peptide, as suggested earlier^2^. In this case the RUST ratio of individual amino acids may not provide an accurate representation of the nascent peptide effect on ribosome movement. Therefore, we measured RUST ratios for peptide fragments (10 residues) with particular physicochemical properties (number of positive charges, net charge and number of hydrophobic amino acids) (Supplementary Fig. 10). Under high positive charge we observed deviations for the distributions of these physicochemical properties in the datasets. However, it is not clear whether they are caused by their direct effects on decoding rates.

We also examined whether tripeptides could affect ribosome velocity differently than may be expected from their individual components. We detect such synergetic effects by comparing the RUST values for tripeptides to what would be expected from independent RUST values of corresponding residues using the standard score (Z-score). We carried out this analysis for adjacent amino acids only and thus explored synergetic effects for 464,000 tripeptides (20 × 20 × 20 residues × 58 positions). Approximately 0.2% (~1,000) of the tripeptides were found to have a standard score greater than 4 (*S_ijk_*>4 or *S_ijk_*< -4) in any individual dataset. These synergistic interactions were found to occur mostly near the decoding center or at the reads termini. They also had a relatively small influence with the majority of interactions having less than a 2 fold change between observed and expected values (Supplementary Fig. 11c). In the ‘Andreev’ dataset the motifs that displayed positive synergetic effects (slower than expected) were overrepresented with Proline. This is a poor substrate for peptide bond formation (see Supplementary Fig. 11a for examples) and therefore a good *a priori* candidate for such synergistic effects. However, there was poor convergence between the results obtained from the thirty datasets, overall 7,854 examples of synergistic interaction were found with the majority (5,850) of candidates found only in a single dataset (Supplementary Fig. 11d).

### Accurate prediction of experimental footprint densities

We proceeded to test whether we can reconstruct ribosome densities using RUST ratios obtained for codon positions relative to the decoding center. Figure 4a shows the comparison of experimental densities to predicted densities based on RUST ratios for the A-site codon or 12 codons comprising the ribosome footprint and the codons immediately adjacent to them. Predictions made based only on the A-site RUST values correlate with the real profiles (Pearson’s *r*=0.451, Spearman’s *r*=0.503 for Andreev et al dataset^27^). The incorporation of RUST ratios for all codon sites in the footprint improves the predictive power even further, with an average Pearson’s r=0.62. These values may improve further with increased sequencing depth. Note that this is not an example of overfitting of a model, as the RUST metafootprint profile is relatively unaffected if it is obtained from a subset of genes (Supplementary Fig. 12) different from which are used to evaluate the profiles. We also compared the profiles to those obtained from another ribo-seq sample of the same study. This had an average Pearson’s r of 0.78, the difference between the samples probably reflects the stochastic nature of RNA-seq. The ability to predict ribosome profiles was replicated for other datasets, with an average Pearson’s correlation coefficient greater than 0.5 in 16 of 30^th^ datasets. The accuracy of these predictions support our earlier findings of a limited influence of the nascent peptide, mRNA structure or synergistic effects on read density. Figure 4b shows comparison of predicted ribosome profiles with experimental profiles for five mRNAs with different degrees of correlation. It is clear from the example shown that the poor correlation is a result of technical artefacts in the data, rather than poor prediction.

**Figure 4.**
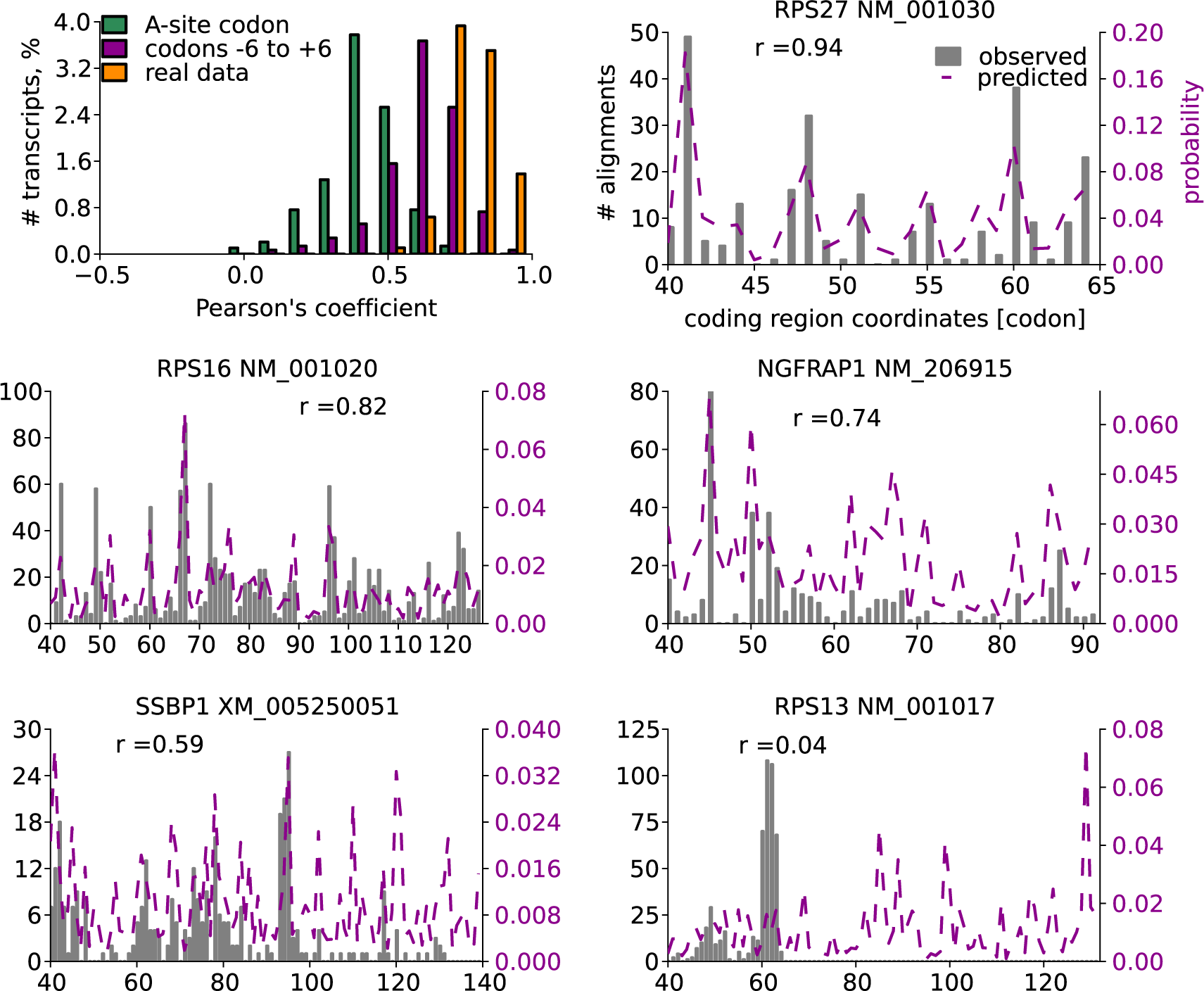
RUST accurately predict experimental footprint densities. Top left panel shows distributions of Pearson’s correlation coefficients for experimental and predicted footprint densities (green and violet) for individual transcripts as well as correlation between two experimental ribo-seq datasets obtained under the same protocol (orange). Correlations were measured only for coding regions of highly expressed transcripts from 120 nucleotides downstream of the start codons to 60 nucleotides upstream of the stop codons. The other panels show experimental (solid grey) and predicted (based on RUST values for the codons -6 to +6 relative to A-site) ribosome densities (broken purple) for five transcripts, corresponding to the 1^st^, 10^th^, 100^th^’, 500^th^ and 714^th^ strongest correlations, the Pearson’s correlation coefficients are indicated. Results displayed are for ‘Andreev’ dataset.

### Comparison of the datasets

We designed a web site http://lapti.ucc.ie/rust, that provides detailed characteristics (metafootprint analysis, RUST ratios, triplet periodicity, etc.) of each dataset explored in this study, an example for an individual dataset is shown in Figure 5. It also hosts executable scripts to implement the RUST analysis.

**Figure 5.**
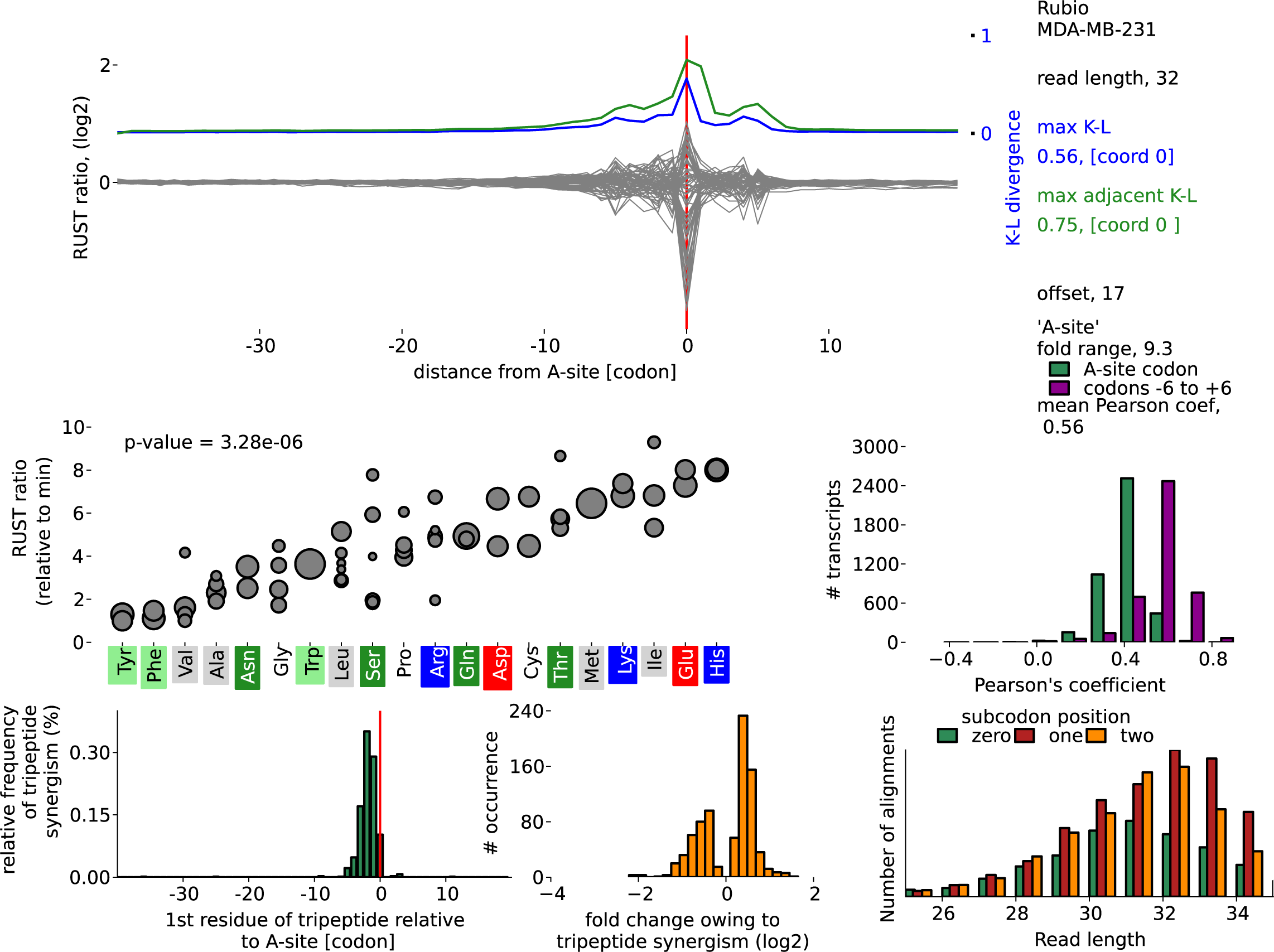
An example of information provided for each dataset with RUST Software. RUST metafootprint analysis for codons is shown at the top. Blue indicates Kulback-Leibler (K-L) divergence for individual codons and green for two adjacent codons. Zero coordinate corresponds to the inferred position of the A-site. Middle left plot shows RUST ratios (Y axis) for individual codons ordered by encoding amino acids at the axis X, the size of the disc indicates the codon usage. Middle right plot illustrates the ability of RUST parameters to reconstruct experimental densities (green – based on identity of the A-site codons and violet is based on all codons within footprints). Bottom left plot shows locations of observed synergistic effects between amino acids that affect decoding rates, the bottom middle plot shows their strength and the bottom right plot illustrates the triplet periodicity signal for footprints of different lengths. This information is available for each dataset used in this study at http://lapti.ucc.ie/rust/.

### Discussion

Here, we described a simple computational technique RUST for the characterization of ribosome profiling data based on a simple smoothing transformation of ribosome density profiles into a binary function. Using simulated data we show that this technique is robust in the presence of sporadic heterogeneous noise (modeled as extra high density and missing data) and outperforms previous methods. Using experimental data, we show that the characteristics of ribosome profiling data extracted with RUST can explain much of the variation observed in experimental ribosome footprint densities.

We applied this technique to thirty publicly available ribo-seq datasets (obtained from yeast, mammalian cultured cells and tissues) and uncovered substantial variability among them in sequence features that determine footprint frequencies at individual locations. The most similar datasets are those obtained with cycloheximide pre-treatments of yeast cells and no or minimal variations in protocols used. For the datasets obtained in mammalian systems we found substantial variation that is likely to be related to the timing of cycloheximide treatments as well as conditions of buffers used for lysis and nuclease digestion. The position specificity of sequencing biases (they affect the boundaries of ribosome footprints) enabled us to determine their relative impact on composition of footprints in individual datasets.

Our simulations suggest that potential uncharacterized artefacts of the computational analysis of ribo-seq data are unlikely to be a major cause in the current difficulties for the determination of the true ribosome decoding rates. However, it appears that all current approaches including RUST overestimate the dwell time of quickly decoding determinants of elongation (codons in the case of simulations). A number of attempts were made to supersede CN approach. Surprisingly, in this study we find that for many applications, such as the analysis of the enrichment rate of individual codons, the simplest variant CN>0 provides surprisingly accurate results. In our simulation we found that it was only marginally worse than RUST irrespective of the relationship between codon usage and dwell times. For real data CN>0 provided results broadly similar to that obtained with RUST, except that the noise reductions achieved with RUST is counterbalanced with a lower signal, (Supplementary Fig. 13). It is likely that the superiority of CN>0 normalization over CN>1 is due to larger volume of data used. While it seems reasonable to filter out lowly expressed genes prior to the analysis because their individual ribosome profiles are unrealistic representations of the real ribosome density, collectively these profiles produce a statistically reliable signal and their analysis is highly informative.

The RUST approach maximizes the chances that detected signal is real in two ways. On one hand it is based on gathering information from all transcriptome coordinates increasing the chance that the signal is not arising due to stochastic reasons. The benefit of this becomes greatest when examining the influence of relatively infrequent determinants, such as certain dipeptides, tripeptides. On the other hand, by reducing the impact of each individual site, RUST ensures that a signal is not a product of a rare outlier (whether due to technical or biological reasons). The smoothing achieved by RUST could also be applied to other high-throughput methods that are characterized by the presence of heterogeneous noise. In this work, for example, we were able to detect that sequencing reads that form RNA secondary structures are underrepresented not only in ribosome profiling data, but also in mRNA-seq data. Thus RUST could have a broader impact if adopted.

The conversion of regular profile to a binary profile leads to an unavoidable loss of information. The approach is therefore “blind” to individual special cases where infrequent motifs may pause the ribosome for a long period. This, however, can be used to identify such special cases by looking for large discrepancies between the densities in the real data and in simulations based on parameters extracted with RUST. This application, however, is challenged by the presence of technical artefacts as illustrated in Figure 5.

We illustrated the applicability of RUST for measurement of mRNA features that impact decoding rates using datasets with lower sequencing bias. The results suggest that sites other than at the decoding center have a relatively minor influence on the decoding rate globally. This observation does not contradict the well characterized pauses modulated by nascent peptide signals and RNA secondary structures at specific locations of individual mRNAs. However, we also showed that in addition to identity of codons in the decoding center of the ribosome, sequences surrounding the ends of footprints are major determinants of footprint densities. The influence of these regions on read density greatly vary among different datasets, in some exceeding that of the sequences in the decoding center. We suggest that this feature could be used for quality assessment of ribosome profiling datasets for the presence of cDNA library construction biases. Crossplatform implementation of RUST is freely available at RiboGalaxy (http://ribogalaxy.ucc.ie).

## Methods

### Ribo-seq datasets used in this study and their processing

The datasets (and SRA repository accession numbers) are summarized in Supplementary Table 1. For simplicity these datasets are indexed in the text using the first author name of the original article. The processing of the reads consisted of clipping the adapter sequence and removal of ribosomal RNA reads followed by the alignment of the mammalian reads to the RefSeq transcriptome^50^ and the yeast reads to the *Saccharomyces cerevisiae* genome (sacCer3 assembly). The weakly updated human RefSeq catalogue was downloaded on the 13^th^ Aug 2014 from the NCBI ftp website ftp://ftp.ncbi.nlm.nih.gov/refseq/H_sapiens/ and the mouse RefSeq catalogue was downloaded on the 18^th^ March 2014 from ftp://ftp.ncbi.nlm.nih.gov/refseq/Mmusculus/. The yeast genome (sacCer3 assembly) and annotation data were downloaded on 13^th^ Aug 2014 from the UCSC genome browser^51^ website, http://hgdownload.soe.ucsc.edu/goldenPath/sacCer3/bigZips/sacCer3.2bit(genome), http://hgdownload.soe.ucsc.edu/goldenPath/sacCer3/database/sgdGene.txt.gz (annotations).

Bowtie version 1.0.0^52^ was used to carry out the alignments. The reads were aligned using Bowtie to the entire human or mouse catalogue with the following parameters (-a, -m 100 -norc). Except where otherwise indicated the reads that mapped unambiguously to a gene (but not necessarily to a single transcript) were brought forward for further analysis. For the yeast datasets reads were aligned to the yeast genome allowing only unambiguous alignments (-a, -m 1).

### Ribo-seq simulation

The simulated alignment data were modelled using real human mRNA sequences obtained from the RefSeq database and with the average transcript read density similar to that of real ribosome profiling data. We simulated the data under the simplistic model where the local decoding rate depends exclusively on the identity of a decoded codon (A-site codon). The number of footprints at each codon position was determined by sampling from the following Poisson probability mass function:

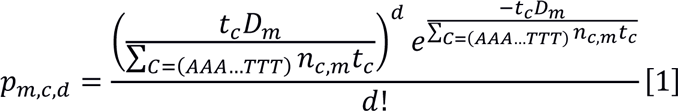

Where *pm,c,d* is the probability of finding *d* number of footprints at a specific location at mRNA *m* at a codon *c* from the set of 61 sense codons *C. Dm* is the total number of footprints aligning to mRNA *m*; *n_c,m_* is the number of codons *c* in the coding region of mRNA *m*; and *t_c_* is the relative dwell time for the codon c. The dwell times *t_c_* for the 61 sense codons were set to span either a ~10 or ~100 fold range with equal increments of 0.15 or 1.5, (the fastest codon was given a score of 1, the slowest was 10.15 or 92.5). To model the noise arising from high density peaks the number of reads at a certain percentage of randomly selected coordinates (irrespective of where it originally contained a mapped read) was substituted with a 3x the value of the highest footprint density for the original simulated profile. The number of codons selected was calculated as a percentage (either 5% or 20%) of the number of codons with a mapped read. To model the absence of mapped reads because of discarding of ambiguous alignments reads were removed from 20% of codons with mapped reads. The selection of codons was carried out using a probability distribution, therefore for individual mRNA density profiles the number of altered codons may differ from 5% to 20%.

The normalisation approaches were used to estimate relative dwell times as described below. The normalised estimated/simulated values obtained for all 61 sense codons were used to produce the boxplots in Figure 1 and Supplementary Figure 2. The normalisation consisted of dividing each of 61 values by their mean to enable comparison between values from datasets and normalisation approaches. The coefficient of determination between the estimated and simulated values was also used as a measure of the accuracy of the approaches.

To explore how the data specific factors (e.g. coverage, sequencing biases) affect performance of different normalization approaches we carried out simulations using Hseih et al^31^ (4 million mapped reads of which 530,051 reads passed selection criteria) and Rubio et al^36^ (61 million mapped reads of which 6,470,387 reads passed selection criteria). The simulations on Figure 1 are based on Rubio et al. data, those in Supplementary Figure 2 were carried out on the Hseih dataset.

### Determining offset to the A-site

An important factor for the analysis described in this work is the application of the correct offset for inferring the position of the A-site codon relative to the footprint 5’ end. This is typically estimated with a metagene profile of either initiating or terminating ribosomes. This may not always allow for a precise estimation of the offset and it is possible that initiating or terminating ribosomes do not protect mRNAs in the same way as elongating ones because of conformational differences, e.g. when release factor eRF1 binds to the ribosome^53^. With the premise that the combined A-site and P-sites should have the greatest influence on decoding rates we set out to estimate the offset using RUST codon metafootprint profiles with the largest K-L divergence at adjacent offsets. We carried out three RUST metafootprint profiles (using the same approach described below) at multiple offsets (usually 16, 17, 18 nucleotides). For these profiles we determined the combined K-L divergence from two adjacent codons. The codon pair in any of the profiles with the largest K-L (that was not at the ends of the reads) was assumed to correspond to the P and A-sites. It was necessary to take the combined K-L divergence from two adjacent sites as in some datasets the divergence of the P-site was greater than that of the A-site. For one of the datasets (with low sequencing bias) we confirmed that the maximal K-L divergence nucleotides corresponded to the A-site offset determined with initiating ribosomes (Supplementary Fig. 14). The offsets used for each dataset are listed in the Supplementary Table 1.

### Normalization approaches

For this analysis the alignment data to the longest coding transcript of every expressed gene were used. Owing to possible atypical translation at the beginning or the end of coding regions, the analysis was carried out on coding regions with the A-site position within 120 nucleotides (40 codons) downstream of the annotated start codon and 60 nucleotides upstream of the annotated stop codon. With exception to one of the ‘Lareau’ datasets the analysis was carried using reads of the predominant length. An offset to the A-site was determined as described earlier. The exclusive selection of reads of one length was necessary to minimise the effect of variation in a distance between footprint ends and the A-site. In this analysis reads were used irrespective of the subcodon position to which they aligned. The exclusive selection of reads that align to a particular subcodon position may further improve the signal. Because of these criteria ~15% of total (non rRNA) mapped reads were used to produce metafootprint profiles (Supplementary Table 1). To check whether exclusion of unambiguously aligned reads had a large influence on the result we repeated the analysis with the unambiguous reads, the obtained results are nearly the same (Supplementary Fig. 15).

The RUST pipeline is described in Supplementary Figure 1. The first step of “RUST phase” is the conversion of ribosome density profile to a binary profile based on whether the number of alignments at each determinant (codon, nucleotide, amino acid) exceeds the gene average. The RUST value at each location *l* is denoted as (*ro_cl_*), *c* stands one of 61 codons (when codons are examined as determinants). For each sequence determinant the expected value *re_cl_* is also obtained. *re_cl_* is obtained by averaging local RUST values across a single coding region. For lowly expressed genes it is expected to be close to 0 and for highly expressed genes it is substantially higher. Normalisation over expected values is carried out to control for the non-random distribution of codons (or other determinants) across the genes with different expression levels. If all codons had the same dwell times, their unnormalized RUST values would be higher for codons that are more frequent in highly expressed genes. This analysis is carried out for all mRNA sequences in the translatome. In order to check for an enrichment of reads at a particular determinant the obtained RUST value is compared to the expected RUST value. To produce a metafootprint profile we used a sliding window approach illustrated in Supplementary Figure 3. For the analysis of codons as a determinant of footprint density the window of 61 codons is moved with a step size of one codon. The center of the window is considered to be the A-site codon. The RUST values are calculated for each codon relative to the A-site and represented in the form of a metafootprint profile. The procedure used for other determinants such as nucleotides, amino acids, peptide properties and RNA secondary structures is conceptually the same.

CN normalisation consisted of an initial normalisation of the individual read density profiles by the average read density specific to individual coding regions. This followed by determination of average normalized values for each of 61 codons across the entire dataset. For the generation of metafootprint profiles average codon values were calculated for specific locations within the sliding window similarly to how it is illustrated for RUST in Supplementary Figure 3. For CN>0 all mRNA transcripts were used while for the CN>1 only coding regions with an average read density > 1 read/nucleotide were used.

We carried out “Ribosome residence time” RRT similar to that described by Gardin et al.^15^ The analyses was carried out independently on windows of 19 codons in length that satisfy the following requirements: (1) greater than 19 aligned reads, (2) less than 3 codons with no alignments and (3) if the codon at the position 10 occurred only once in the window. For each window the decimal fraction of reads aligning to each codon (relative to the total number of reads in the window) was determined. The average obtained for each codon at all 19 codons was then used to produce the metafootprint profile.

As the other normalisation procedures do not use mRNA-seq data, we could not carry out an equitable comparison with the “corrected ribo coverage” analysis^12^. Therefore, instead of using the footprint density normalized by RNA-seq density, we used only footprint densities. We refer to this approach as LMN for logarithmic mean normalisation. Similar to the original approach only coordinates with mapped reads are used and footprint densities are first normalised by the algebraic average read density. The algebraic average of their log_2_ values are then calculated across all coding regions (first term in equation [2] below). Further the average of all 61 codons is calculated (second term in equation [2] below) and subtracted from the codon specific value. The procedure can be summarized in the following equation

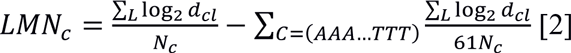

Where *LMNc* is LMN value for the codon c (from a set of 61), *Nc* is the total number of *c* codon occurrences with non 0 footprint densities and *d_cl_* is footprint density for the codon *c* at the location *l* normalized by the average footprint density for the corresponding mRNA. We carried out the analysis on coding regions with an average read density > 1 read/nucleotide.

The “aov” function in R was used to calculate the p-values with ANOVA for assessing statistical significance of the difference between the variation among synonymous codons and variation among all codons at the A-site.

### Kullback-Leibler divergence

The Kulback-Leibler divergence was used to calculate relative entropy in the RUST metafootprint profiles and was calculated as the following

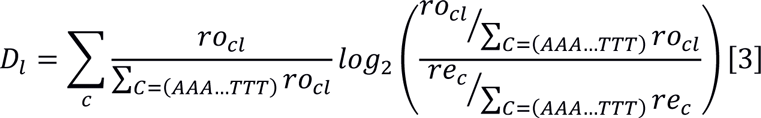

Where *D_l_* is the K-L at location *l*, *ro_cl_* is the observed RUST value for codon *c* at location *l* and *re_c_* is the expected RUST value for codon *c*. The higher the K-L, the less uniform the distribution of RUST values is in the corresponding position. Thus, K-L indicates how much the corresponding position contributes to the abundance of footprints.

### RNA secondary structure analysis

The computational prediction of RNA secondary structure free energy was performed using RNAfold in the ViennaRNA package^49^. Using a sliding window of 80 nucleotides with a step size of 10 nucleotides the minimal free energy for potential RNA secondary structures was estimated across each transcript. For human data the threshold free energy for the most stable RNA secondary structures was found to be-40.1 kcal/mol for the top 1^st^ percentile, -32.8 kcal/mol for the 5^th^ and −29.0 kcal/mol the 10^th^ percentile.

### Amino acid physicochemical properties

In this study Histidine, Lysine, Arginine, were considered to be positively charged. Aspartic acid, Glutamic acid as negatively charged. Alanine, Valine, Isoleucine, Leucine, Methionine, Phenylalanine, Tyrosine, Tryptophan were considered to be hydrophobic.

### Standard score to identify synergistic interactions

To identify synergistic interactions, we compared the difference in fold changes between observed and expected metafootprint profiles. The fold change at each position was normalised to the background fold change as follows

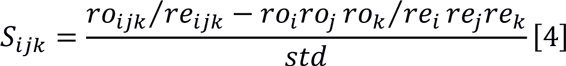

Where *S*_*ijk*_ are synergy indexes for tripeptide *ijk* and *ro/re* are corresponding RUST ratios. *std* is the standard deviation of the differences observed at regions from -40 to + 18 relative to the A-site.

### The comparison of predicted and real footprint densities

When information from all footprint codons, plus two surrounding ones (-6 to +6 relative to the P-site/A-site boundary) was used to model ribo-seq densities, the predicted profile can be represented as a discrete probability density function

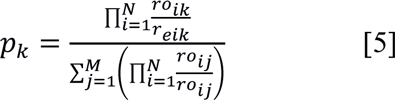

Where *p_k_* is the probability of finding a footprint at the position *k* of the mRNA coding region consisting of *M* codons. *ro_ik_/ro_ik_* is the RUST ratio for the codon at the site *i* (relative to the codon *k*) from the total of *N* sites used. For instance, if RUST ratios of AAA in the P-site and A-site are 0.339 and 1.646 respectively, the expected RUST ratio for di-codon AAA-AAA is 0.557 (0.339*1.646). Instead of di-codons in our simulation the RUST ratio is obtained with 12 codons, this corresponds to the numerator in equation [5]. The denominator corresponds to the sum of RUST ratios across the coding region and remains constant for all codons of each transcript.

The comparison between the expected and experimental profiles was carried out on transcripts with a density greater than 1 read/nucleotide. (Transcripts with a lower density were not used as they have insufficient data to correlate with the predicted profile).

The python package matplotlib^55^ was used to produce the figures.

### Code availability

Supplementary Software is a compressed archive of user friendly executable scripts to run RUST (version 1.2). Its source code is accessible and it includes several implementations of RUST that search for enrichment of codons, amino acids, dipeptides, tripeptides and nucleotides. “rust_synergy” searches for synergistic effects between adjacent amino acids. “rust_predict_profiles” returns a csv file that records the Pearson’s and Spearman’s correlation coefficient between the observed and predicted footprint densities for individual transcripts. “rust_plot_transcript” plots the observed and predicted footprint densities. This (and updated versions in the future) are also available at http://lapti.ucc.ie/rust/. In addition, Supplementary Software includes “RUST_script.py” a 2^nd^ shorter, non-executable version of the RUST implementation on codon enrichment. This script is a pseudocode intended as an explanatory aid for understanding RUST algorithm. RUST is also available via the RUST package at RiboGalaxy^54^ (http://ribogalaxy.ucc.ie).

### Data availability

The NCBI SRA accessions numbers for the datasets processed in this study is listed in Table 1

## Acknowledgments

We would like to thank Audrey Michel for critical reading of the manuscript. We also are grateful to Can Cenik, Justin Gardin, Nicholas Ingolia, Shu-Bin Qian, Noam Stern-Ginossar, Craig Stumpf, and Jonathan Weissman for providing us with the details of experimental protocols that were used to generate the datasets surveyed in this work.

This work was supported by grants from Science Foundation Ireland (12/IA/1335) and the Wellcome Trust (094423) to P.V.B.

## Author Contributions

P.B.F.O’C and P.V.B. conceived the study. P.B.F.O’C developed the method and carried out the data analysis. D.E.A. surveyed ribosomal profiling protocols. All authors participated in interpretation of the data. P.B.F.O’C and P.V.B. wrote the manuscript.

